# When the heart and the brain meet: Brain-heart interactions in feature integration

**DOI:** 10.64898/2026.05.22.726192

**Authors:** María I. Cobos, Clara Alameda, Pedro M. Guerra, Ana B. Chica

## Abstract

In contemporary Cognitive Neuroscience, increasing attention is devoted to brain-body interactions, as an expanding body of literature suggests that processing the external world may not emerge from the brain in isolation but rather from the coordinated contribution of multiple bodily systems. These interactions have been extensively studied in the context of interoception. However, evidence linking them to visual perception remains scarce. To address this gap, the present study reanalyses a previously published dataset from 25 participants to investigate heart-brain interactions during a visual feature integration task. Participants were required to integrate shape and colour features to identify a target while inhibiting distractor-related information. Cardiac and neural activity were simultaneously recorded, enabling the assessment of the heart rate (HR), heart-evoked potentials (HEP), and, albeit seldom reported previously, heart-evoked oscillations (HEO). Pre-stimulus HEP amplitude briefly differed between correctly and incorrectly integrated features. HEO analysis revealed alpha and low beta band modulations before target onset. These effects were absent when the analyses were aligned to external events and exceeded surrogate distributions obtained after disrupting heartbeat–EEG alignment, indicating that they depended on cardiac timing rather than on spontaneous prestimulus EEG differences alone. These findings provide preliminary evidence that HEO dynamics are associated with successful perceptual integration and extend previous work on HEPs from stimulus detection to higher-level perceptual processes. More broadly, they suggest that heartbeat-locked neural activity may influence early brain states linked to perceptual outcomes, aligning with theoretical frameworks proposing an active role for bodily signals in perceptual processing.

**Highlights:**

- A brief pre-stimulus HEP amplitude differs between correct and incorrect feature integration.
- Alpha and low beta HEO increases before correct feature integration.
- Pre-stimulus oscillatory effects disappeared when analyses were time-locked to external events rather than cardiac activity.
- Convergent HEP and HEO findings suggest a role for brain-heart interactions in visual perception

## 1. Introduction

Throughout the history of Cognitive Neuroscience, research on cognitive processes has mainly focused on brain dynamics. However, growing evidence suggests that the brain does not operate in isolation but instead engages in continuous bidirectional communication with the organs of the body (Blanke et al., 2015; Christoff et al., 2011; Craig, 2002; Critchley & Harrison, 2013; Damasio & Carvalho, 2013; Garfinkel & Critchley, 2016; Park & Tallon-Baudry, 2014). This ongoing interaction is of particular interest because it shapes how we perceive, process, and interact with the world around us.

Building on this brain-body perspective, contemporary theories of consciousness are currently grappling with a range of emerging challenges, one of which is understanding the role that the organism itself plays in the process of conscious perception (Cleeremans et al., 2025). This approach is not new; for example, Damasio (2010) proposed that conscious experience emerges from the interaction between the mind and the organism, modulated by emotional and homeostatic processes. According to this view, consciousness results from the functional integration of neural maps monitoring the body’s internal physiological state with the maps representing changes in the external environment (Damasio, 2010; Damasio & Damasio, 2023). Similarly, Park & Tallon-Baudry’s (2014) “neural subjective frame” extends this view, proposing that reporting a conscious experience (“I have seen the stimulus”) requires a subjective sense of self constructed from the integration of exteroceptive (e.g., vision) and interoceptive (e.g., heartbeats) information. This integration would form what is known as the first-person perspective, with heart-evoked potential (HEP) being one of its proposed neural signatures (Loescher et al., 2025; Park & Tallon-Baudry, 2014; Tallon-Baudry et al., 2018). In a pioneering study, Park et al. (2014) found that HEP amplitude increased when a visual stimulus was consciously detected (as compared to missed stimuli), even before the stimulus appeared. Additionally, HEP modulations have been linked to bodily self-consciousness (Park et al., 2016, 2018), with a greater HEP amplitude during asynchronous versus synchronous bodily illusions. While these findings suggest that HEPs are associated with different facets of conscious experience, perceptual HEP studies have primarily focused on conscious detection. However, conscious perception involves not only access to sensory information but also the integration of information into coherent perceptual representations (Kouider et al., 2007, 2010; Lamme & Roelfsema, 2000; Storm et al., 2024). Accordingly, feature integration paradigms provide a useful framework for examining conscious processes beyond stimulus detection alone. Together, these theoretical and empirical accounts highlight the cardiovascular system as a unique and tractable window onto brain-body interactions.

Investigating heart-brain interactions requires the use of complementary indices capturing different aspects of this bidirectional relationship. For example, heart rate (HR) provides a direct measure of cardiac activity, while also reflecting the influence of the brain through autonomic efferent pathways. At the same time, the HEP, recorded at the brain level, reflects neural responses modulated by cardiac signals, enabling the simultaneous study of cardiac and neural contributions to cognitive processes. Both indices have been widely used as markers of brain-heart dynamics in the study of emotion (Couto et al., 2015; Critchley et al., 2005; Fukushima et al., 2011; Gentsch et al., 2019; Lucas et al., 2019), and stress (Schulz et al., 2013; Thayer et al., 2012), as well as in attention (Fourcade et al., 2024; Luft & Bhattacharya, 2015; Sforza et al., 2000) and interoception (Canales-Johnson et al., 2015; Garfinkel et al., 2015; Pollatos et al., 2005; Pollatos & Schandry, 2004). Conversely, their role in perception has been less explored, particularly in the case of HEPs. While HR studies have reported larger cardiac decelerations associated with response preparation (Jennings et al., 2009; Jennings & van der Molen, 2005; van der Veen et al., 2004), conscious perception (Cobos et al., 2019; Motyka et al., 2019; Park et al., 2014), and error detection (Cobos et al., 2026; Di Gregorio et al., 2024; Skora, Livermore, Nisini, et al., 2022; Wessel et al., 2011), evidence linking HEPs to perceptual processes remains comparatively scarce.

Apart from evoked responses, as with HEP, growing interest is emerging in the relationship between cardiac-evoked activity and oscillatory brain dynamics. In particular, Luft & Bhattacharya (2015) reported a positive correlation between HEP amplitude and alpha power across different arousal levels. Interestingly, Grosselin et al. (2018) revealed alpha and beta-band modulations evoked by cardiac activity during wakefulness. This approach was one of the first to introduce the possibility that the heart may also evoke oscillatory responses (heart-evoked oscillations: HEO). Lechinger et al. (2015) similarly reported positive correlations between HR and alpha power during awake vs. sleep states. More recently, heartbeat-modulated oscillations have been associated with both HEP activity and interoceptive performance, providing further evidence that cardiac-related oscillatory dynamics may have behavioural relevance (Zaccaro et al., 2026). Although sparse, these studies are relevant because they point toward an additional way to explore heart-brain interactions associated with brain oscillations.

Given the scarcity of studies examining heart-brain interactions in perceptual processing and the relevance of identifying markers of the first-person perspective proposed by the *neural subjective frame* (Park & Tallon-Baudry, 2014), the present study aimed to characterize brain-heart interactions during feature integration. To that aim, we performed a reanalysis of a previously published dataset (Cobos et al. 2023), leveraging a feature integration paradigm as an established approach to studying perceptual integration processes involved in conscious perception (de Gardelle & Kouider, 2009; Kouider et al., 2007, 2010). Specifically, participants performed a feature integration task adapted from Esterman et al. (2004, 2007), in which a string of characters with different shape and color features was briefly presented. Responses involved either correct or incorrect feature integration, with the latter giving rise to perceptual illusions. In addition, the role of top-down expectancy was explored by introducing an unexpected stimulus feature at the end of the experiment, which only some participants noticed.

Brain-heart dynamics were characterized using HR, HEP, and HEO. Following the pre-registered plan (https://osf.io/3zv5r), we hypothesized a deceleration-acceleration pattern in HR (Skora, Livermore, Nisini, et al., 2022; Skora, Livermore, & Roelofs, 2022), replicating previous findings (Cobos et al., 2026). At neural level, we expected increased HEP amplitudes for correct versus incorrect feature integration (Park et al., 2014) and explored potential post-target differences without directional predictions. We also predicted larger HEP amplitudes in aware compared to unaware participants. However, the awareness factor was not significant in any of the pre registered analyses. Because these analyses did not alter the interpretation of the main findings, they are reported in full in the Supplementary Material section 6 to improve the readability of the main text while maintaining transparency. Furthermore, we predicted pre-target HEP amplitudes to correlate with cardiac acceleration during response emission, as well as positive associations between HEP amplitude and both behavioral performance (proportion of correct integrations) and alpha band oscillatory activity (Luft & Bhattacharya, 2015).

## 2. Methods

In the present study, we conducted a reanalysis of the data previously published by Cobos et al. (2023). Accordingly, we provide a summary of the experimental methods. For further details or any additional technical information, please consult the original work by the authors (Cobos et al., 2023).

### 2.1 Participants

Thirty healthy volunteers (21 females; M = 24 years, SD = 2.87) participated in the study in exchange for 10€/h. All participants reported normal or corrected-to-normal vision and color perception, and no prior experience with the task. Written informed consent was obtained, and participants were informed of their right to withdraw at any time. The study was approved by the CEIM/CEI Granada’s Biomedicine Ethics Research Committee and conducted in accordance with the 1964 Declaration of Helsinki.

The primary statistical analyses were performed using cluster-based permutation tests (Maris & Oostenveld, 2007), which do not permit a direct parametric estimation of statistical power. To provide a standard quantification of the design’s sensitivity, a sensitivity analysis was conducted in G.Power 3.1 (Faul et al., 2007), modelling a paired-samples t-test equivalent to the contrast between the two experimental conditions (correct and incorrect feature integration). With N = 27, α = .05 (two-tailed), and a medium-large power of 0.80, the minimum detectable effect size was dz = 0.56, indicating that the available sample was sufficiently sensitive to detect effects of moderate magnitude.

### 2.2 Stimuli, apparatus and procedure

The experimental display started with a central fixation point (white plus sign, 0.47° × 0.47°) on a black background (see Figure 1 for a detailed description). Subsequently, a number (1-9, excluding 5) was presented 0.95° above fixation. The peripheral stimulus consisted of a horizontal string of four characters printed in Arial font, initially sized at 3.8° × 1.05°, and presented 5.7° to the left or right of fixation. This string contained two white external flankers (“S” or “8”) and two inner letters (“L” and “O”). The inner letters were always in different colors, chosen from red, blue, or green. The “L” served as the target and the “O” as the distractor. A mask (“&&&&”, subtending the same size as the string of characters) followed the character string.

**Figure 1:**
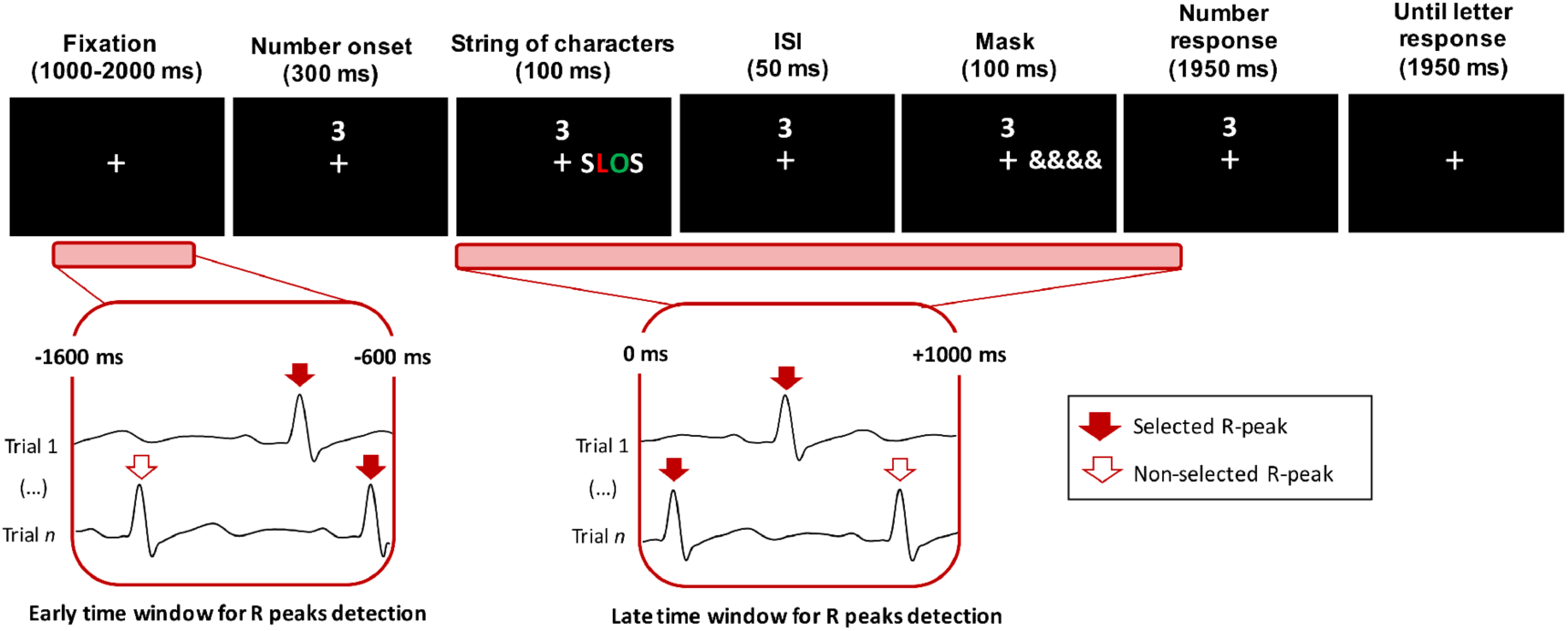
(Top): Sequence and timing of events in a given trial. Participants first responded to the number task, reporting if the number was larger or smaller than 5, as fast and as accurately as possible. After this, participants responded to the color of the letter L as accurately as possible. (Bottom): Early and Late HEP windows, durations, and selected R-peaks. Last R peak before target onset (left side), and first R peak after target onset (right side).

Stimulus presentation and behavioral data collection were controlled by E-Prime 2.0 software. Stimuli were displayed on a 24” LCD monitor (Benq BL2405HT, 1920 × 1080 pixels, 60 Hz refresh rate), with participants seated approximately 60 cm away.

The experiment comprised 14 blocks of 48 trials each, preceded by a titration procedure. This titration adjusted the peripheral stimulus size in 0.3° × 0.1° increments until a participant achieved ∼70% accuracy, ensuring a sufficient rate of illusions for analyses. Participants first responded to the number task (right-hand mouse response), indicating if the number was smaller or larger than 5. Here, both speed and accuracy were emphasized. Then, participants had to indicate the color of the letter (red/blue/green) using the left-hand keys “z”, “x”, or “c”. The space bar could be pressed if participants could not identify the color of the target (unseen trial). This response prioritized accuracy over speed. The responses were classified as hits (correct target color), illusions (reporting the color of the “O” distractor), errors (incorrect color, not displayed), or unseen.

To investigate the role of top-down expectancy in feature integration, the experimental design transitioned from ten “expected blocks” to four “unexpected blocks”. During the initial expected blocks, the target letter was always colored (red, blue, or green), establishing a strong perceptual set. Unknown to the participants, in the final four blocks, a white target was introduced on 50% of the trials. Because the response keys were only designated for the three primary colors, these trials had no correct response (therefore, only illusions, errors or unseen responses were recorded).

At the end of the session, the experimenter conducted a brief structured interview to assess whether participants were aware of the white letter or not.

### 2.3 Recording and preprocessing

#### 2.3.1 EEG acquisition and preprocessing

EEG was recorded using a 64-channel actiCAP system (Brain Products) at 1000 Hz, with impedances kept below 5 kΩ and online reference at FCz. EOG activity was monitored using two electrodes (TP9–TP10) placed at the outer canthi.

EEG preprocessing followed the pipeline described in Cobos et al. (2023). Only trials where participants correctly responded to the number task and responded to the string of characters were included in the analysis. Six preprocessing and artifact-rejection steps were applied. 1) Data was segmented into long epochs (4000 ms) to avoid edge effects. 2) The power-line artifact at 50Hz and the harmonics (100, 150 Hz) were reduced using spectrum interpolation (Leske & Dalal, 2019), 3) Baseline correction was applied from -2000 to 0 ms to improve EEG trace visualization. 4) EEG data was visually inspected trial-by-trial and participant-by-participant. Trials containing artifacts were manually removed. Trials with blinks or eye movements occurring 600 ms before or 800 ms after number onset were also excluded. In total, 11% of trials were discarded on average (SD=6.5). 5) Independent Component Analysis (Fieldtrip “runica” algorithm) was used to remove remaining blinks. 6) An average of 2.37 channels (SD=2.12) was interpolated using neighboring electrodes. Two participants were excluded because their rejected-trial count exceeded two standard deviations above the mean, and another participant was removed for having nine bad channels. Finally, EOG electrodes were removed from the analysis, and EEG data were re-reference to the common average. Full methodological details can be found in Cobos et al. (2023).

#### 2.3.2 ECG acquisition and preprocessing

ECG was recorded using a Biopac MP150 system connected to a PC running AcqKnowledge software (version 3.9.1.6). HR was derived from the electrocardiogram (ECG) recorded at Lead II using a Biopac MEC110C module. The ECG signal was sampled at 1000 Hz and band-pass filtered between 0.5 and 35 Hz during acquisition.

Subsequently, a digital FIR high-pass filter was used within Acqknowledge software to eliminate frequencies lower than 2.5 Hz (Blackman window with a 61dB roll-off). ECGlabRR software (Vicente et al., 2013) was employed to derive HR from the ECG data. Initially, each cardiac cycle was assessed by measuring the interval, in milliseconds, between consecutive R waves, which was then converted into beats per minute. Following this, Kardia software (Perakakis et al., 2010) was used to calculate the weighted average of the HR for each trial every 100 ms over a 2500 ms interval that was time-locked to the fixation point. The HR differential scores were determined by substracting the average HR activity that occurred during the 1000 ms immediately preceding the onset of the fixation point (baseline).

#### 2.3.3 HEP preprocessing

HEPs were computed from continuous EEG data time-locked to R-peaks extracted from the ECG signal. EEG and ECG recordings were temporally aligned offline using the experimental triggers to ensure precise correspondence between cardiac events and stimulus timing before peak detection.

For R-peak detection, R-peak latencies were first identified in the continuous ECG signal using the MATLAB HEPlab toolbox (Perakakis, 2019) and then visually inspected to ensure accuracy. Subsequently, the R-peaks occurring within the two-time windows of interest were selected (for a similar procedure, see Park et al. (2014) : 1) early time window (i.e., before number presentation), to explore preparatory components, from -1600 ms to -600 ms relative to target onset, and 2) late time window (i.e., after target presentation) to explore feature integration components, from 0 to 1000 ms after target onset (see Figure 1, bottom). In the event that more than one R-peak was found within the defined windows, the second R-peak was selected in the early time window, and the first R-peak in the case of the late time window (see Figure 1, filled red arrows). This procedure ensured that the generated epochs were closer to the time points of interest (i.e., immediately before and immediately after target presentation). Importantly, in the preparatory window, the 200– 600 ms post–R-peak analysis interval did not overlap with any task-related stimulus, ensuring that the measured activity reflected ongoing preparatory brain-heart dynamics rather than stimulus-evoked responses. Furthermore, a control analysis confirmed that the selected R-peaks were similarly distributed relative to target onset in hits and illusions (t (24) = 0.28, p = .782, Cohen’s dz = 0.06; see Supplementary Figure S4). EEG epochs were extracted from −200 to 600 ms relative to the selected R-peak and baseline-corrected using the −200 to 0 ms interval. Due to behavioral performance differences, the number of trials differed between conditions. In the preparatory window, the mean number of trials per participant was 254.2 (SD = 66.93) for hits and 114.64 (SD = 54.08) for illusions. In the post-target window, hits: M = 254.08, SD = 66.88; illusions: M = 115.00, SD = 54.04.

#### 2.3.4 Exploratory HEO preprocessing

HEO signal was extracted by applying a time-frequency (TF) analysis implemented in FieldTrip (Oostenveld et al., 2010). HEO epochs were extracted at 1000 ms before and after R-peak selection. To minimize edge effects in the TF analysis, each epoch was extended by 750 ms on both sides with mirror padding, yielding a final segment of 3000 ms. The extracted epochs were longer due to padding, but the wavelet transform was computed within a -1000 to 1000 ms window, ensuring analysis was based on non-padded data. Power was calculated for each trial using the Wavelet approach, with frequencies ranging from 2 to 32Hz in 0.25Hz steps (Candia-Rivera et al., 2021; Cannard et al., 2024; Grosselin et al., 2018). The number of cycles increased linearly from 3 at low frequencies (<20Hz) to 7 at high frequencies, thus adapting the temporal window to optimize resolution across the spectrum. In both cases, the signal was analyzed in 25ms steps. The resulting TF maps were normalized at the participant level by calculating the relative change from baseline (−300 to −200 ms locked to R-peak) (Candia-Rivera et al., 2021; Cannard et al., 2024; Grosselin et al., 2018).

### 2.4 General analysis plan

#### 2.4.1 HR analyses

HR analyses were restricted to the expected blocks (480 trials) for the hits and illusions conditions when the number task was responded to correctly.

Outliers were removed in those instances where the mean of all the experimental conditions was greater or smaller than 2.5 standard deviations (N=1). Another participant was excluded from HR data due to ECG recording failure (N=1).

HR was analyzed with linear mixed-effects models (LME) using *lme4* (Bates et al., 2014) in RStudio. The dependent variable was HR (beats/min). Fixed effects included Trial type (hit vs. illusion), Time (25 levels sampled every 100 ms), and their two-way interaction (HR measurement ∼ Trialtype * HR_Level).

To account for the hierarchical structure of the data, several candidate random-effects structures were tested while keeping the fixed-effects structure constant. The simplest model included a random intercept for Participant (1∣Participant). A second model added a random intercept for Trial type nested within Participant to capture trial-wise heterogeneity (1∣Participant) + (1∣Participant:Trialtype). A third model additionally included a random slope for Trial type across Participants, allowing the hit–illusion effect to vary between participants (1+Trialtype∣Participant) + (1∣Participant:Trialtype). A fourth model further included a random slope for Trial type across Participants (1 + Trialtype|Participants) + (1 + Number trial|Participants) + (1|Participants:Number trial), in order to model potential linear trends across the experimental session; this model also retained the random intercept for trials nested within participants. The optimal random-effects structure was found using a data-driven model comparison by running the *anova* function of the stats package of R (Chambers & Hastie, 1992). The lowest AIC and BIC parameters indicated the fittest model to choose the random factor. When AIC and BIC yielded different lowest values, the simplest model among them was selected. Then, the *Anova* function of the *car package* was used to explore the effects (Fox et al., 2022). The effect size for LME was informed by the eta square (η2), extracted by the *eta_square* function of the sjstats R package (Lüdecke et al., 2025).

#### 2.4.2 HEP analyses

From the total sample, 27 participants provided valid EEG data after preprocessing. Additionally, two participants were eliminated due to ECG recording failure (N = 1, reported above) and the inability to detect R-peaks (N = 1). Therefore, the final sample for the HEP analyses consisted of 25 participants.

Statistical comparisons between hits and illusions HEP amplitude were conducted using a non-parametric cluster-based permutation test (Maris & Oostenveld, 2007) implemented in FieldTrip (Oostenveld et al., 2010)^1^. Analyses were restricted to the 200–600 ms interval following the R-peak as the literature supports this timing for identifying the HEP effect (Coll et al., 2021). In addition, this window was selected to minimize contamination from cardiac field artifacts immediately following the R-peak, as recommended by previous work (Park et al., 2014). A two-tailed dependent-samples t-test was used as the sample-level statistic, with a cluster-forming threshold of p < .05. Clusters were formed across neighbouring electrodes, requiring a minimum of two spatial neighbours, and cluster mass was evaluated using the “maxsum” statistic. The null distribution was estimated using 5,000 Monte-Carlo permutations, yielding cluster-level p-values that control the family-wise error rate.

#### 2.4.3 Exploratory HEO analyses in early time window

The statistical procedure was identical to the cluster-based permutation test (Maris & Oostenveld, 2007) described in section 2.4.2, but adapted to time–frequency analysis. Power was computed in FieldTrip (Oostenveld et al., 2010), and hit vs. illusion differences were tested within the 200–600 ms post-R-peak interval and the 4–30 Hz frequency range. Statistical parameters were equal to those in section 2.4.2. The initial cluster-based permutation test was conducted across the entire TF space (4–30 Hz, all electrodes, 200– 600 ms post-R-peak) to identify the specific frequency range contributing to any significant effect (see Supplementary table S2). Subsequent analyses were then restricted to the frequency interval revealed by this data-driven cluster, which was examined within canonical bands (theta: 4–7 Hz; alpha: 8–12 Hz; low beta: 13–20 Hz; high beta/gamma: 20– 30 Hz).

## 3. Results

### 3.1 HR results

HR analysis showed a main effect of Time (X^2^(24) = 32360.48; p<.001; η^2^ = 0.09), revealing a deceleration pattern (see Figure 2). No other main effect or interactions reached significance (all ps>.87).

**Figure 2:**
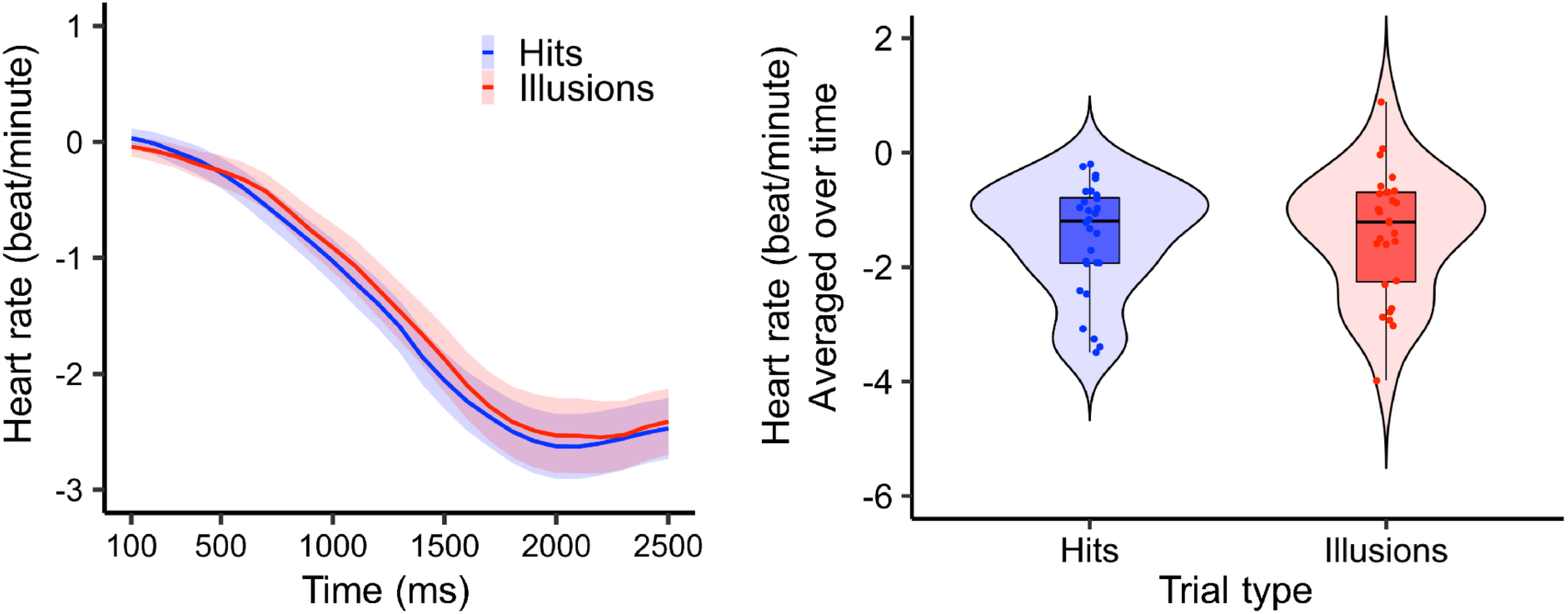
Left panel: Changes in HR (relative to baseline) over time for hits and illusions, time-locked to fixation onset. Target appearance occurred from 1300 to 1400 ms. The main effect of time was significant. The standard error of the mean (SEM) for each time point is represented by a shaded area of color. Right Panel: HR response (averaged over time) for each experimental condition of Trial type. The main effect of Trial Type was not significant. The maximum and minimum values per condition are represented in the whiskers of the box plots. The Interquartile range (IQR) is displayed in the boxes by portraying the lower quartile, median, and upper quartile. The colored dots represent mean values for each participant. The violin shape depicts the kernel density estimation of the distribution of participant-level means, where wider sections indicate higher data density.

### 3.2 HEP results

Cluster-based permutation testing of the early HEP (R-peaks immediately preceding target onset) revealed a cluster-level effect differentiating hits and illusions between 486 and 506 ms post–R-peak (cluster-level p = .027; 5,000 permutations). This effect was observed over parieto-occipital electrodes (O1, Oz, O2, P4, P8, PO7, PO3, POz, PO4, PO8, and P6; black dots in Figure 3), where HEP amplitude was more negative for hits than for illusions. No additional effects were detected in the early window (all ps > .068).

**Figure 3.**
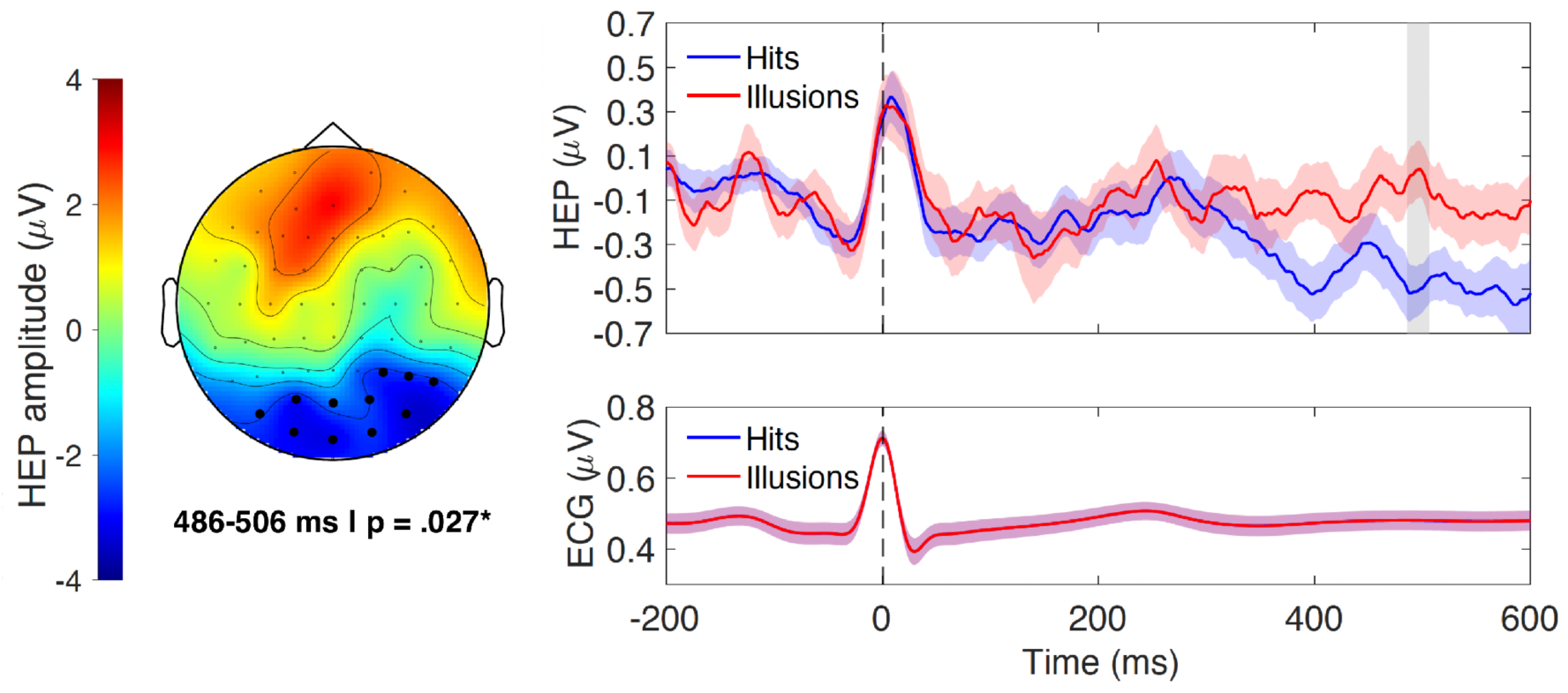
Cluster-based permutation testing identified a significant difference between hits and illusions in the 486–506 ms interval (gray shaded area) post R-peak (cluster-level p = .027), over parieto-occipital electrodes. Left: Topographical distribution of t-values averaged across the significant interval; black dots indicate electrodes contributing to the cluster-level statistic. Right: Grand-average HEP waveforms (mean ± SEM) extracted from these electrodes, computed across all participants and all trials within each condition (top), smoothed using a 20 ms moving window for visualization purposes, with the grey area indicating the temporal extent contributing to the cluster statistic. The corresponding mean ECG waveform for each condition is shown below.

In the late window (R-peaks following target onset), no reliable differences were observed between hits and illusions (all ps> .105).

Additionally, to verify that potential HEP differences were not attributable to peripheral cardiac differences, ECG waveforms were compared between conditions after alignment. No differences were observed in R-peak amplitude or morphology between hits and illusions in either time window (see Figure 3, bottom right), indicating that any subsequent effects reflect differences in neural activity rather than cardiac signal variability.

### 3.3 HEO results in early time window

Cluster-based permutation tests across the full frequency range (4-30 Hz) revealed two significant positive effects in the early time window (see Table S2 in the Supplementary material). The completed cardiac-locked TF map for each experimental condition and their differences is shown at the top of Figure 4. The first effect reflected differences between hits and illusions (p=.017; 5,000 permutations) in the 325-600 ms post-R-peak interval and the frequency range of 6.7-21 Hz, as illustrated by the shaded area in the upper panel of the TF map in Figure 4-c. This effect was observed in a set of 23 centro-parieto-occipital electrodes (C2, C4, C6, CP3, CP5, CP6, FC2, FC4, FC6, O1, O2, Oz, P3, P5, P6, P7, P8, PO3, PO4, PO7, PO8, TP7, TP8; see the upper plot in Figure 4-c). The second effect was observed between 200-300 ms and a frequency range of 8.3-21 Hz (p=.040; 5,000 permutations), as shown by the shaded area in the lower panel of the TF map in Figure 4-c. This effect was observed in a set of 21 centro-parieto-occipital electrodes (C2, C4, CP4, CP6, FC2, FC4, O1, O2, Oz, P3, P4, P5, P6, P7, P8, PO3, PO4, PO7, PO8, POz, TP8; see the bottom topoplot in Figure 4-c,). For both significant effects, greater cardiac-locked oscillatory power was observed for hits than illusions.

**Figure 4:**
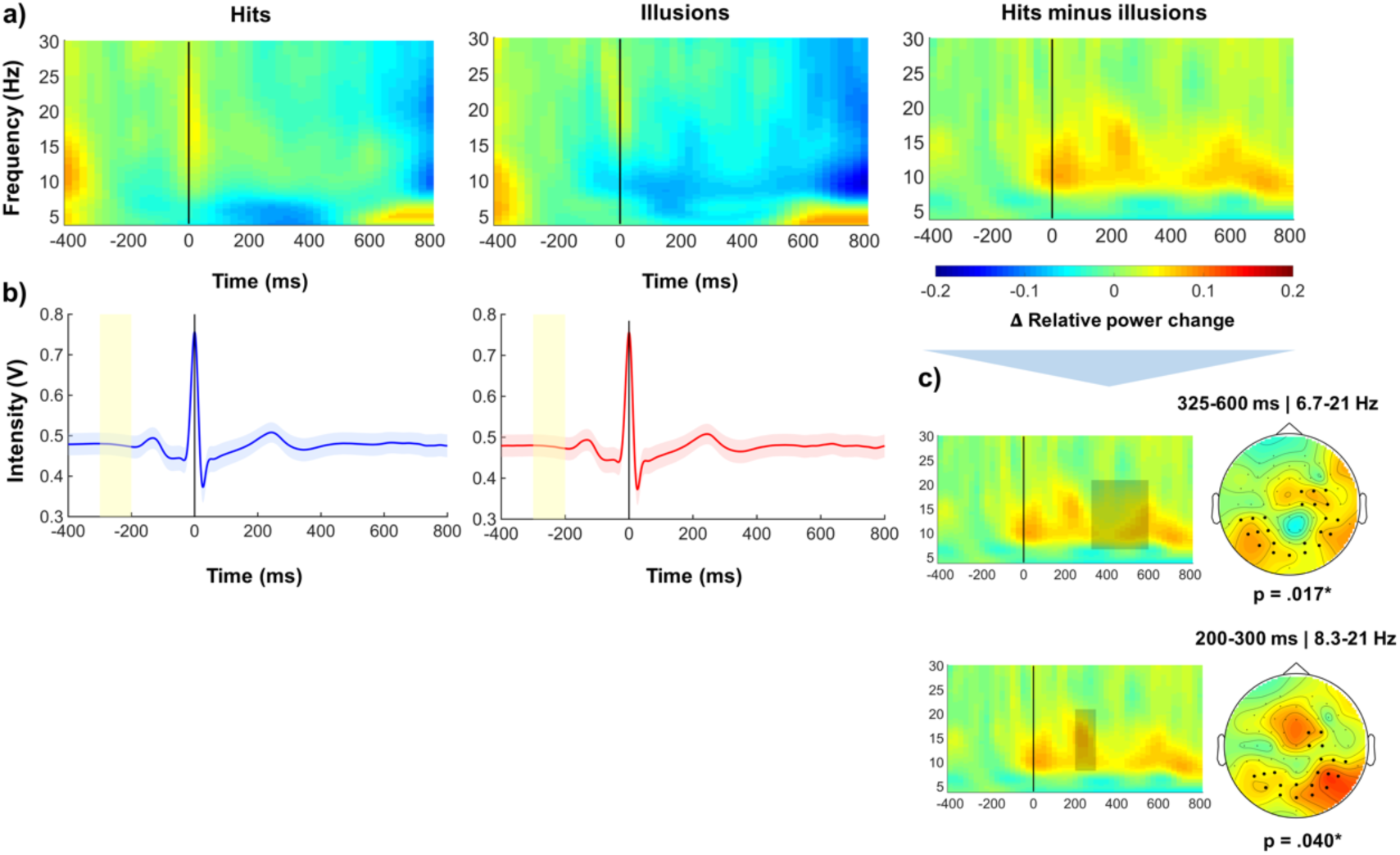
Cluster-based permutation analysis representation across the completed frequency range (4-30 Hz). The analyzed window was from 200 to 600 ms after R-peak. a) The figure shows the TF map for hits, illusions, and the difference between hits and illusions (from left to right). b) In the hits and illusions conditions, the participant’s average of the ECG signal is represented below the corresponding TF map. The colored shaded area indicates the SEM; the yellow shadow shows the baseline time. c) TF map for each significant cluster (top cluster-level p = .017; bottom cluster-level p = .040), along with the corresponding topographic distribution of the averaged t-values within the significant TF intervals; black dots indicate electrodes contributing to the cluster-level statistic; the color bar is the same for the TF map; the asterisk represents the significant results (p < .05*). To improve visualization, wavelet-related missing values (NaNs) in the TF maps were masked and interpolated using a nearest-neighbor approach, ensuring a continuous TF representation without altering the underlying estimates.

The cluster-based permutation test for the specific range of frequencies indicated that theta power (4-7 Hz) did not reach significance (p=.330; 5,000 permutations). However, alpha power (8-12 Hz) exhibited significant differences between hits and illusions over centro-parieto-occipital electrodes (C2, C4, C6, CP5, CP6, FC2, FC4, FC6, O1, O2, Oz, P5, P6, P7, P8, PO3, PO4, PO7, PO8, TP7, TP8). Specifically, greater cardiac-locked alpha power was observed for hits compared to illusions (p=.024; 5,000 permutations). These differences were observed between 326 and 600 ms after the R-peak (see Figure 5-a).

**Figure 5:**
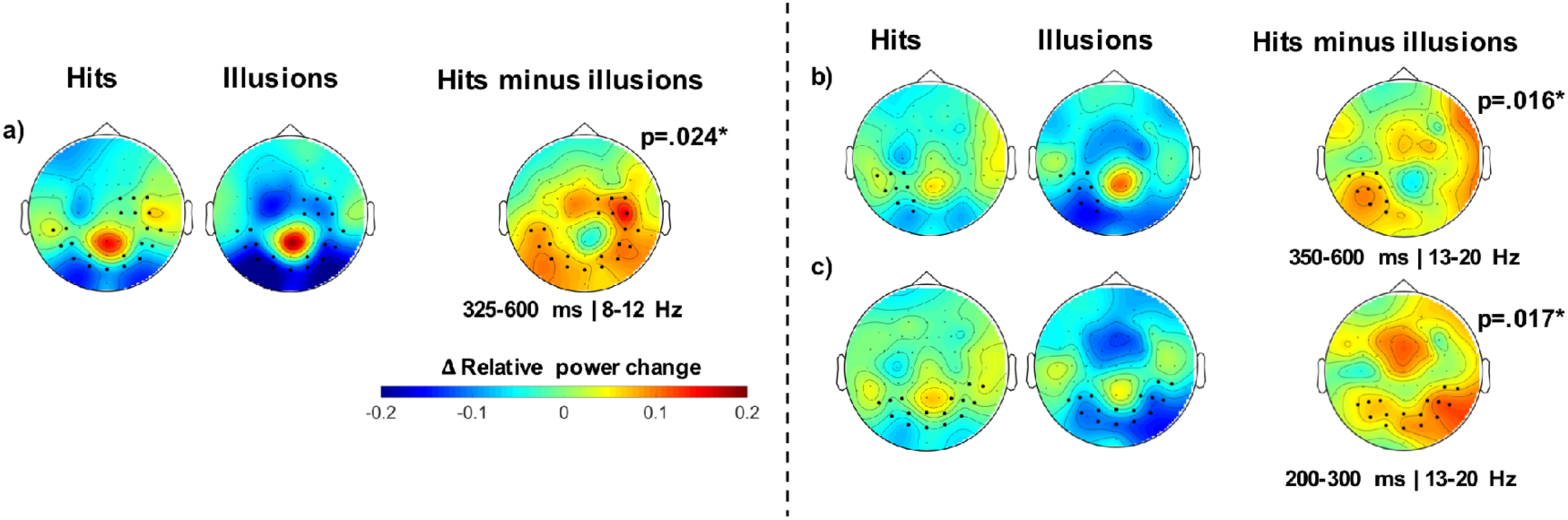
Cluster-based permutation significant results in the specific frequency ranges. a) Topographical distribution of t-values averaged across the significant time intervals in alpha (8-12 Hz) for hits, illusions, and the difference between hits and illusions. a) Significant differences between hits and illusions were observed in the alpha range (8-12 Hz) from 325 to 600 ms post R-peak (cluster-level p = .024), over centro-parieto-occipital electrodes. b) Significant differences in the low beta range (13-20 Hz) were found between 350 and 600 ms post R-peak (p=.016) over left parieto-occipital electrodes. c) A further significant low beta effect was observed earlier, between 200 and 300 ms post R-peak (p=.017), also over parieto-occipital electrodes. Black dots indicate electrodes contributing to the cluster-level statistic; the asterisk represents the significant results (p < .05*).

Within the low beta power (13-21 Hz) two significant effects were observed (see Figure 5 b-c). The first effect, with a left parieto-occipital distribution (CP3, CP5, O1, P3, P5, P7, PO3, PO7, TP7), showed greater cardiac-locked beta power for hits than for illusions (p=.016; 5,000 permutations) during the 350-600 ms post-R-peak interval (see Figure 5-b). A second parieto-occipital effect (CP6, O1, O2, Oz, P3, P4, P5, P6, P7, P8, PO3, PO4, PO7, PO8, POz, TP8), also revealed greater cardiac-locked beta power for hits relative to illusions (p=.017; 5,000 permutations) from 200 to 300 ms (see Figure 5-c).

### 3.4 Post hoc stimulus-locked TF control analysis

To verify that the power effects reported in the HEO analyses did not simply reflect classical preparatory alpha modulations unrelated to cardiac processing, a post hoc analysis was conducted on the same temporal windows without using ECG-derived information to define the temporal reference of the analysis . This verification was aimed at determining whether differences observed in the HEP and HEO analyses related to Trial Type (hits vs. illusions) would also emerge when neural oscillations were locked to the presentation of external stimulation. The oscillatory dynamics were analysed using the general wavelet-based approach described in Section 2.4.3. The same preprocessed EEG trials were re-epoched relative to fixation onset (comparable to the early time window in Figure 1), and target onset (comparable to the late time window in Figure 1). TF estimates were computed from -1500 to 1500 ms relative to each task event. Both event-locked analyses used the same 300 ms baseline period (-500 to -200 ms relative to fixation onset; -800 to -500 relative to target onset) No additional trial rejection was applied; therefore, the number of trials and the hit/illusion imbalance were identical to those used in the heartbeat-locked analyses. For each event-locked analysis, statistical testing covered a 1000 ms period starting from the respective time locking event.

The results of the full frequency range time-locked to the fixation point did not reach statistical significance (all ps>.106). However, when the analysis was time-locked to the target, significant differences were observed (p=.033; 5,000 permutations) from 775 to 1000 ms. This effect involved a set of 26 electrodes (AF3, AF7, AFz, C1, C2, C3, C5, CP2, CP4, CP6, CPz, Cz, F1, F3, F5, FC1, FC3, FC5, Fz, O2, P4, P6, P8, PO4, PO8, TP8) in the frequency range of 7.5-14.8 Hz. The results revealed a greater power decrease for hits compared to illusions (see Supplementary Figure S3). Alpha frequency (8-12 Hz) exhibited significant differences (p=.017; 5,000 permutations) from 775 to 1000 ms in a set of 24 electrodes (AF3, AF7, AFz, C1, C2, C3, C5, CP2, CP4, CP6, CPz, Cz, F1, F3, F5, FC1, FC3, FC5, Fz, P4, P6, P8, PO8, TP8), with a greater decrement in alpha for hits than illusions. The analyses of the remaining frequencies did not reach significance (all ps>.073).

### 3.5 Additional surrogate-heartbeat control analyses

To assess whether the reported HEP and HEO effects depended on the temporal correspondence between heartbeat timing and neural activity, we performed additional surrogate-heartbeat control analyses following recent methodological recommendations (Steinfath et al., 2025, 2026). In these analyses, heartbeat timing was randomly reassigned across trials while preserving condition-related EEG activity. Both the HEP (p = .042) and HEO (p = .035) effects exceeded the surrogate distributions obtained after disrupting heartbeat–EEG alignment. These findings indicate that both effects depended on the temporal alignment between heartbeat timing and EEG activity, rather than being explained by spontaneous prestimulus EEG differences between experimental conditions alone. Full methodological details and surrogate distributions are provided in Supplementary Sections 4.2 and 4.3 (Supplementary Figures S5 and S6).

### 3.6 Correlation analyses

According to the preregistration and the potential relationship between HEP amplitude, behavioral performance, and HR outcomes, different correlations were conducted using the function *cor.test* from the stats package in Rstudio. Indexes were computed for each modality to assess whether Trial type differences were related across physiological and behavioral measures. For the early HEP, an amplitude index was obtained by subtracting the amplitude for illusions from the amplitude for hits. Likewise, a behavioral index was calculated as the difference between the proportion of hits and the proportion of illusions. A first correlation examined the association between early HEP and behavioral indexes. A second analysis focused on HR measures: the minimum value of the HR deceleration and the minimum HR value before target presentation (up to 1300 ms) were extracted, and corresponding hit minus illusion indexes were computed. Both HR indexes were then correlated with the early-HEP index. Spearman’s rank correlation coefficient was used as a robust, non-parametric measure of association.

Neither the correlation between the HEP and the behavioral indices (Spearman’s rho=0.211; p = .310), nor the correlation with the minimum point of cardiac deceleration (Spearman’s rho=-0.072; p = .743), nor the correlation with the minimum HR value before target presentation (Spearman’s rho=-0.077; p = .726) reached significance. Importantly, HEP and alpha power correlation had been planned. However, we realized that the pre-target window examined here does not overlap with the time window of early alpha power reported in Cobos et al. (2023), making such a comparison temporally inappropriate. Exploratory trial-level analyses were additionally conducted to examine whether prestimulus HEP amplitude and HEO power predicted subsequent perceptual outcome. Greater trial-to-trial HEP negativity within participants was associated with a lower probability of reporting an illusion (β = −0.078, OR = 0.925, 95% CI [0.884, 0.968], p = .00074), whereas analogous HEO analyses yielded effects in the same direction but did not reach statistical significance (all ps>.093; see Supplementary Section 5)

## 4. Discussion

The present study investigated whether brain-heart interactions are related to visual feature integration, a process closely linked to subjective aspects of conscious perception (de Gardelle & Kouider, 2009; Kouider et al., 2007, 2010). Our findings show that pre-stimulus cardiac-related neural activity may be associated with subsequent perceptual experience. Specifically, correct feature integration was preceded by distinct HEP responses and HEO dynamics compared to incorrect feature integration. Crucially, these effects were observed in heartbeat-locked alpha and low beta activity before target onset and disappeared when the analyses were instead aligned to external events, a finding further supported by surrogate-heartbeat control analyses, providing preliminary evidence that HEO dynamics may play a role in successful feature integration. We also observed a temporally restricted pre-stimulus HEP effect comparable to that reported by Park et al. (2014), extending the investigation of brain-heart interactions from conscious detection to feature integration. Importantly, these effects emerged independently of HR changes, suggesting that correct perception may depend more strongly on the functional interaction between cardiac and cortical activity than on peripheral cardiac or brain dynamics alone.

Unlike previous studies reporting HR differences associated with visual perception accuracy (Cobos et al., 2019, 2026; Park et al., 2014), HR did not differ between correct and incorrect feature integration in the present study. One important methodological difference may explain these contrasting findings: previous studies included warning signals before stimulus presentation, either on every trial (Park et al., 2014) or in some trials (Cobos et al., 2019, 2026), whereas no alerting cues were presented here. Since warning signals reliably modulate HR (Jennings & van der Molen, 2005; Lacey & Lacey, 1978; Skora, Livermore, & Roelofs, 2022; Vila et al., 2007), earlier HR effects may reflect preparatory cardiac modulation that influences stimulus detection at the conscious threshold or feature integration. The absence of HR differences in the present study, therefore, suggests that successful feature integration can occur without detectable changes in peripheral cardiac activity.

In line with Park et al. (2014), our results revealed a brief pre-stimulus HEP modulation, supporting the involvement of heart-brain interactions in visual perception. However, whereas Park et al. (2014) contrasted detected and undetected stimuli, the present study examined correct and incorrect feature integration. Thus, rather than extending previous findings on conscious detection alone, our results suggest that pre-stimulus cardiac-related activity may also be associated with the successful integration of perceptual information into a coherent representation. Supporting this view, somatosensory perception studies by Al and colleagues (2020, 2021) have also reported a larger pre-stimulus HEP for hits compared to misses. Alongside this evidence, Park et al. (2016, 2018) reported increased HEP amplitudes in a bodily illusion paradigm when visuo-tactile stimulation was synchronous compared to asynchronous (illusory perception). This enhanced HEP activity was associated with improved body self-awareness. Taken together, this body of evidence raises the possibility that increased HEP amplitude may reflect a more coherent alignment between cardiac and cortical signals during perceptual processing. However, the specific cognitive processes underlying this relationship remain unclear. Successful feature integration depends on multiple interacting mechanisms, including attentional mechanisms (Briand, 1998; Cobos & Chica, 2022; Cohen & Rafal, 1991; Grubb et al., 2013; Montaser-Kouhsari & Rajimehr, 2005; Prinzmetal et al., 1986; Yeshurun & Rashal, 2010), distractor suppression (Capilla et al., 2014; Klimesch et al., 2007; Lange et al., 2014; Min & Herrmann, 2007; Schroeder et al., 2018), and feature binding (Herrmann et al., 2004; Phillips & Takeda, 2009; Singer, 2013; Tallon-Baudry, 2009), and the current HEP results do not allow direct associations with specific cognitive perceptual processes. Moreover, despite growing evidence linking HEP and HEO activity to perceptual and interoceptive processes, the broader literature has not yet established robust associations between specific HEP modulations and particular cognitive-perceptual functions (Coll et al., 2021; Steinfath et al., 2026). Our data suggest that pre-stimulus HEP modulations may support the perceptual system toward more efficient feature integration, thereby increasing the likelihood of veridical perception.

While the present findings and previous work emphasize the relevance of HEPs as markers of heart-brain interactions, evoked responses may capture only part of the influence of cardiac signals on cortical processing. HEP amplitude is closely related to ongoing alpha activity and varies with arousal states (Luft & Bhattacharya, 2015), and cardiac-locked alpha has been associated with global brain states such as wakefulness (Grosselin et al., 2018), pointing to oscillatory dynamics as a key pathway for cardiac influences on brain function. However, HEOs have rarely been examined in active cognition. Building on this framework, we found distinct pre-stimulus HEO modulations in the alpha and low beta bands for correct and incorrect feature integration. In particular, alpha power was higher for correct than incorrect feature integration during the pre-target window. This finding aligns with previous evidence linking alpha dynamics to perceptual accuracy and conscious perception (Di Gregorio et al., 2022; Kienitz et al., 2022; VanRullen, 2016). Alpha oscillations have been proposed to act as a temporal sampling mechanism shaping perceptual sensitivity (Haegens et al., 2014; Hülsdünker et al., 2016; VanRullen, 2016) consistent with findings that faster alpha frequency predicts greater perceptual accuracy, while alpha amplitude and phase influence sensory sensitivity and metacognition (Battaglini et al., 2020; Samaha & Postle, 2015; Zhou et al., 2021). Accordingly, our results suggest that HEO modulation of alpha power may reflect a physiological pathway through which the cardiac signals could influence optimal sampling states, aligning with evidence that alpha dynamics are state-dependent and flexibly adjust to optimize information processing (Mierau et al., 2017). Thus, brain-heart interactions may be involved in stabilizing alpha oscillations that underlie rhythmic perceptual cycles (VanRullen, 2016), thereby improving feature integration.

Low beta power was also higher for correct than for incorrect feature integration during pre-target window. Beta activity has traditionally been associated with motor processes (Engel & Fries, 2010). Interestingly, a recent study has highlighted heartbeat-locked beta activity as a potential marker of cardiac signal processing during interoceptive attention (Pultsina et al., 2026), suggesting that beta oscillations may constitute a relevant component of heart-brain interactions. However, greater pre-stimulus beta power predicts successful visuospatial perception in visual crowding (Ronconi & Bellacosa Marotti, 2017), and can causally enhance performance when low beta stimulation is applied to parietal regions (Battaglini et al., 2019; Di Dona et al., 2024). Further evidence indicates that parietal beta-band modulations influence visual stimulus identification (Capotosto et al., 2009). The enhanced pre-stimulus low beta power observed here occurs before stimulus onset, thus preceding motor preparation or response, and suggesting a perceptual rather than motor role. Recent accounts extend beta function to include top-down feedback processing and long-range communication within perceptual networks, particularly via parietal regions of the dorsal stream (Bastos et al., 2015; Di Dona & Ronconi, 2023; Spitzer & Haegens, 2017). This interpretation aligns with source localization studies showing low beta rhythms in parietooccipital regions (Capilla et al., 2022) and with HEP activity in the right inferior parietal lobe (rIPL) that is stronger for hits than misses (Park et al., 2014). Thus, our pre-stimulus low beta enhancement may reflect integrated brain-heart interactions which could facilitate correct perception, potentially involving parietal regions.

Interestingly, alpha and low-beta frequencies overlapped in timing. Several studies have linked desynchronization in these bands to perceptual fidelity (Griffiths et al., 2019), memory retrieval (Michelmann et al., 2016), visual perception (Pfurtscheller et al., 1996), and the processing of event-related information across cortical modules (Jensen & Mazaheri, 2010; Klimesch, 2012). However, these findings are based on target-locked oscillations. Our results show for the first time that heartbeat-locked alpha and beta modulations in the pre-stimulus interval are associated with visual feature integration. To determine whether these effects depended on cardiac time-locking, we conducted post hoc control analyses replicating the same TF pipeline while removing cardiac information and time-locking activity to external stimuli (fixation point and target). These analyses revealed that pre-target frequency modulations disappeared, whereas post-target results replicated the expected alpha decrease for hits compared to illusions, consistent with Cobos et al. (2023). Although preparatory oscillatory effects have been previously reported (Borra et al., 2023; Ergenoglu et al., 2004; Parés-Pujolràs et al., 2023; Wutz et al., 2018), our findings show that during the preparatory phase, alpha and low beta activity was locked to cardiac signals rather than to the visual stimulus, whereas during target processing, oscillations were locked to the stimulus rather than to cardiac activity. These findings are particularly relevant because they suggest that, before the target appears, preparatory activity is preferentially aligned with internal bodily signals, whereas during target processing it becomes more strongly aligned with external sensory input. Importantly, this does not imply that heartbeat-related modulations are absent during task processing, or that irrelevant stimuli cannot influence neural activity. Rather, it suggests that during periods in which visual stimuli are not relevant (i.e., fixation periods), neural responses are synchronized with cardiac activity, whereas during periods in which visual stimuli are relevant (i.e., target presentation), neural responses are synchronized with the processing of those stimuli. These findings suggest that the brain could have not only a temporal sampling mechanism that shapes perceptual sensitivity (Haegens et al., 2014; Hülsdünker et al., 2016; VanRullen, 2016), but also a dynamic switching mechanism that fluctuates between synchronizing with internal bodily signals and external sensory input. Such flexibility could allow the brain to anticipate salient changes both in the environment and within the organism, thereby facilitating the alignment of internal and external information, potentially contributing to responses to behaviorally relevant events.

The identification of HEO dynamics in the alpha and low beta bands represents a key contribution of this study, extending previous research beyond HEP responses and suggesting that temporal alignment between cardiac signals and ongoing neural oscillations may be involved in visual perception. Importantly, these findings open the possibility of interpreting HEO as a novel marker of the body’s involvement on perception, potentially reflecting aspects of the first-person perspective or, more conservatively, the functional role of cardiac signals in conscious perceptual processing (Azzalini et al., 2019; Cleeremans & Tallon-Baudry, 2022; Damasio, 2010; Damasio & Damasio, 2023; Park & Tallon-Baudry, 2014; Tallon-Baudry et al., 2018). Within this framework, heart-brain synchronization may be linked to more perceptually accurate outcomes, supporting the view that bodily signals may play an active role in, rather than merely accompany, correct perception.

Despite the innovative insights provided here, several limitations must be acknowledged. First, the current study was based on a reanalysis of a previously published dataset and the HEO analysis was exploratory. Consequently, these findings should be considered preliminary, albeit promising, until replicated in an independent dataset focused on testing HEO activity. Another limitation concerns the unequal number of hit and illusion trials, an inherent feature of paradigms designed to produce perceptual illusions. Although our preregistered plan included balancing trial numbers through repeated subsampling, this approach yielded highly variable cluster solutions across iterations (see section 1.3 of the Supplementary material). We therefore conducted the primary analyses using all available trials to maximize estimation stability. Importantly, the cluster most consistently identified across the subsampling procedure corresponded to the main effect reported here (see section 1.2 of the Supplementary material). The current study did not provide subjective measures, such as confidence scales, to assess participants’ certainty about their performance. Such data would provide greater insight into the more subjective aspects of conscious processing (Fleming & Lau, 2014; Jimenez et al., 2025). Additionally, no significant participant-level correlations between the HEP and behavioral measures were observed. However, exploratory trial-level analyses suggested an association between prestimulus HEP fluctuations and subsequent perceptual outcome, whereas analogous HEO analyses showed effects in the same direction but did not reach statistical significance. Notably, this result should nevertheless be interpreted cautiously until replicated in a planned independent dataset. Likewise, correlations between the HEP and HR may have been absent because our results lack a clear HR acceleration pattern associated with response emission. Finally, the present findings are correlational in nature and therefore do not allow causal conclusions regarding the role of brain-heart interactions in feature integration. Moreover, although the task targeted feature integration, the observed heart-brain interactions effects may also reflect other perceptual processes. Future research using causal approaches will be necessary to clarify the specific mechanisms underlying HEP and HEO effects.

Taken together, our findings suggest that heartbeat-locked neural activity may reflect early cortical states potentially influencing the efficiency of feature integration. Our results are consistent with the neural subjective frame proposal (Park & Tallon-Baudry, 2014; Tallon-Baudry et al., 2018) and extend previous work by identifying HEO modulations in alpha and low-beta bands before successful feature integration. Furthermore, these novel HEO findings represent a particularly novel contribution, extending previous research beyond HEP responses and suggesting that brain-heart interactions are associated with perceptual processing through oscillatory activity. Although the specific cognitive processes underlying these HEP and HEO modulations remain to be determined, our study provides converging evidence that heart-brain interaction plays a broader role in the perceptual domain than previously recognized, beyond its well-established relevance for interoception and emotion.

## Authorship contribution

**Ana B. Chica:** conceptualization, funding acquisition, methodology, project administration, resources, supervision, validation, writing – review and editing.

**Clara Alameda:** conceptualization, data curation, formal analysis, methodology, software, validation, visualization, writing – review and editing.

**María I. Cobos:** conceptualization, data curation, formal analysis, funding acquisition, investigation, methodology, resources, software, validation, visualization, writing-original draft, writing – review and editing.

**Pedro M. Guerra:** conceptualization, data curation, funding acquisition, methodology, resources, software, validation, writing – review and editing.

## Declaration of Generative AI and AI-assisted technologies in the writing process

During the preparation of this work, the author(s) used Microsoft Copilot in order to assist with grammar checking and to suggest clearer phrasing to improve readability. The manuscript was entirely written by the author(s), and the tool was used solely to refine the language and ensure clarity of expression. After using this tool/service, the author(s) reviewed and edited the content as needed and take(s) full responsibility for the content of the published article.

## Acknowledges

This work was supported by research grant PID2023-152001NB-I00 funded by MICIU/AEI/ 10.13039/501100011033; ERDF/EU to Ana B. Chica and by research grant PID2020-119549GB-I00 funded by MCIN/AEI/10.13039/50100011033 to Pedro M. Guerra and by research grant CEX2023-001312-M funded by MICIU/AEI/10.13039/501100011033 to María I. Cobos.

The authors would like to thank Dr. Luis F. Ciria for his valuable feedback, insightful comments, and helpful suggestions that contributed to the development of this work.

## Supplementary information

Supplementary information related to this article can be found in the OSF link: https://osf.io/3zv5r

## Data availability

All EEG and ECG data, analysis files, and home-made Matlab and Rstudio scripts can be accessed through the open-access platform Open Science Framework (OSF). Please use the following link to access all materials:

https://osf.io/3zv5r

Project doi: https://doi.org/10.17605/OSF.IO/3ZV5R

## Footnotes

1 ***Deviation from preregistration***. The preregistered analysis plan included a parametric bootstrapping procedure (1,000 repetitions) to balance unequal trial numbers between conditions by random subsampling. In the present dataset, this approach yielded unstable cluster solutions across repetitions, with substantial variability depending on the selected trial subset. To provide a more stable and statistically interpretable estimate of condition differences, the primary analyses were conducted using all available trials. Full subsampling results are reported in the Supplementary Materials for transparency.

