## Supplementary_material for "When the heart and the brain meet: Brain-heart interactions in feature integration": SUPPLEMENTARY MATERIAL_HeartEvokedStudy_R1_24_10_26_clean.pdf

### 1. Pre-registered HEP analyses: subsampling-based parametric bootstrapping

#### *1.1 Procedure*

To address the imbalance in trial numbers between Hits and Illusions, we implemented the preregistered subsampling-based parametric bootstrapping procedure. For each participant, trials from the condition with the larger number of observations (typically Hits) were randomly subsampled to match the number of Illusion trials. Cluster-based permutation testing (as described in the main Methods) was then performed at the group level. This procedure was repeated 1,000 times, each iteration using a different random subset of trials. Due to computational constraints, repetitions were executed in blocks of 50 and subsequently aggregated. For each iteration, the cluster-based outputs (*stat*, *prob*, and *mask*) were stored. These fields were averaged across repetitions to obtain a consensus representation of the results. In this framework, *mask* reflects the proportion of iterations in which each channel–time point survived cluster-level correction. Consensus clusters were defined as spatiotemporally contiguous points exceeding a recurrence threshold of ≥5% across iterations. For each cluster, we extracted its latency range, spatial extent, number of significant channel–time samples, and recurrence metrics (mean, median, and maximum frequency). Cluster polarity was determined from the mean sign of the corresponding *stat* values.

#### *1.2 HEP bootstrapping results*

No cluster survived correction in more than 50% of the 1,000 subsampled iterations. Thus, no spatiotemporal pattern demonstrated high stability across random subsets of trials. However, three consensus clusters exceeded the ≥5% recurrence threshold. These clusters are summarized in Supplementary Table S1 and visualized in Supplementary Figure S2 (left column: topographies; right column: corresponding HEP waveforms).

The most recurrent cluster (Cluster 1) was a posterior negative cluster spanning 0.485–0.514 s post–R-peak (14 electrodes; mean recurrence frequency = 12.2%; maximum frequency = 16.8%). A second posterior negative cluster (0.389–0.408 s; 17 electrodes) showed a mean recurrence frequency of 7.4%. A smaller frontal positive cluster (0.399–0.411 s; 8 electrodes) showed a mean recurrence frequency of 5.3%.

**Supplementary Table S1**

| Idx | Valence | nCh | Time (s) | Size (points) | MeanFreq | MedianFreq | MaxFreq | Electrodes |
|---|---|---|---|---|---|---|---|---|
| 1 | Negative | 14 | 0.485–0.514 | 244 | 0.122 | 0.137 | 0.168 | O1, Oz, O2, P4, P8, CP6, P5, PO7, PO3, POz, PO4, PO8, P6, CP4 |
| 2 | Negative | 17 | 0.389–0.408 | 190 | 0.074 | 0.076 | 0.096 | Pz, P3, P7, O1, Oz, O2, P4, P8, P1, P5, PO7, PO3, POz, PO4, PO8, … |
| 3 | Positive | 8 | 0.399–0.411 | 50 | 0.053 | 0.054 | 0.055 | Fz, F3, F4, AF3, AFz, F1, AF4, F2 |

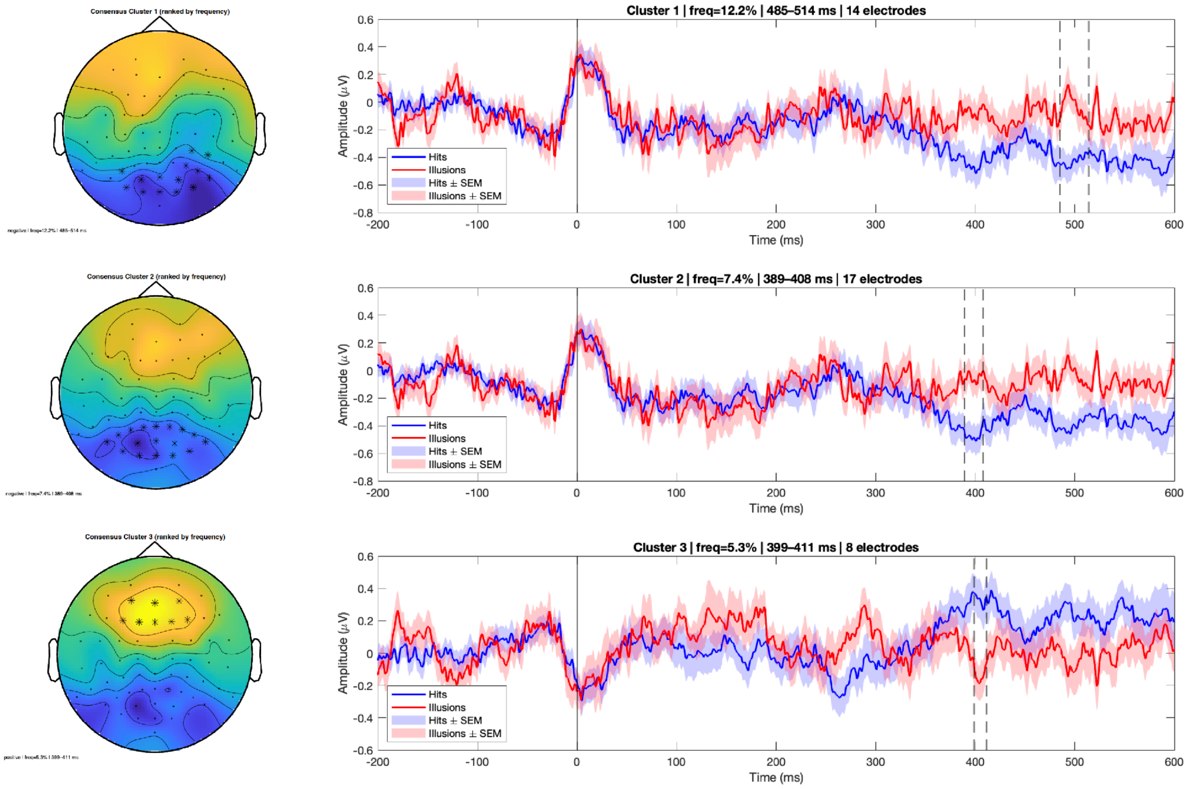


**Supplementary Figure S2. Consensus clusters from the subsampling-based parametric bootstrapping analysis.** Left: Scalp topographies of the three consensus clusters identified across 1,000 subsampled cluster-based permutation iterations, ranked by mean recurrence frequency. Stars indicate electrodes contributing to each cluster. Warmer colors represent positive t-values (Hits > Illusions), and cooler colors represent negative t-values (Illusions > Hits), averaged across iterations within the cluster time window. Right: Grand-average HEP waveforms (± SEM) for Hits (blue) and Illusions (red) extracted from the electrodes comprising each cluster. Vertical dashed lines indicate the temporal extent of each consensus cluster. The grey shaded region marks the analyzed 200–600 ms post–R-peak interval. Cluster recurrence frequencies (freq), latency ranges, and number of contributing electrodes are indicated in each panel.

Importantly, although the spatial distribution and latencies of these clusters were broadly consistent with the main hypothesis and prior literature (Coll et al., 2021) on preparatory HEP effects, their recurrence across subsamples was low. The specific configuration of significant clusters varied considerably between iterations, indicating that statistical outcomes were highly dependent on the particular subset of trials selected in each repetition.

#### *1.3 Limitations of the subsampling approach*

The subsampling procedure substantially reduced the number of trials in the Hit condition, discarding a large proportion of available data in each iteration. This reduction in signal-to-noise ratio, combined with cluster-level multiple-comparison correction, likely increased the probability of Type II errors, particularly for small-to-moderate effects. Moreover, combining repeated subsampling (1,000 iterations) with cluster-based correction imposes a double layer of conservativeness: an effect must not only survive cluster correction within an iteration, but also recur consistently across many random subsets of trials. In the present dataset, this approach led to low reproducibility of cluster solutions and limited interpretability of the consensus maps.

For these reasons, and to provide a more stable estimate of condition differences while preserving statistical power, the primary analyses reported in the main manuscript were conducted using all available trials within a single cluster-based permutation framework. The subsampling results are reported here for transparency and completeness.

### 2. Supplementary HEO data

#### *2.1 HEO table results*

| Cluster valence | Time (ms) | Frequency range (Hz) | Number of channels | Mean t value | Peak t value | p value |
|---|---|---|---|---|---|---|
| Positive | 325-600 ms | 6.7-21.0 Hz | 23 | 3.0261 | 5.3792 | *0.0172 |
| Positive | 200-300 ms | 8.3-21.0 Hz | 21 | 2.8247 | 4.4009 | *0.0404 |
| Positive | 200-275 ms | 11.0-16.7 Hz | 14 | 2.8606 | 4.6604 | 0.1358 |
| Positive | 300-350 ms | 21.7-24.7 Hz | 4 | 2.4592 | 3.3622 | 0.5159 |
| Positive | 400 ms | 15.0-16.7 Hz | 4 | 3.0937 | 4.2447 | 0.6553 |
| Positive | 425 ms | 29.3-30.0 Hz | 4 | 2.7042 | 3.0145 | 0.7461 |

**Supplementary Table S2**: The results of the cluster-based permutation test were analyzed within a time window from 200 to 600 ms post R-peak and over the frequency range of 4-30 Hz. Two significant positive clusters were demonstrated; the asterisk represents the significant results (p < .05*).

### 3. Supplementary TF data without cardiac activity

#### *3.1 TF table results*

| Cluster valence | Time (ms) | Frequency range (Hz) | Number of channels | Mean t value | Peak t value | p value |
|---|---|---|---|---|---|---|
| Positive | 525 ms | 28.8-30.0 Hz | 3 | 2.2896 | 2.5771 | 0.9612 |
| Negative | 775-1000 ms | 7.5-14.8 Hz | 26 | -2.6918 | -3.9614 | * 0.0332 |
| Negative | 550-775 ms | 12.0-16.2 Hz | 10 | -2.9119 | -4.2968 | 0.1808 |
| Negative | 0-150 ms | 10.2-14.5 Hz | 7 | -2.6141 | -3.7295 | 0.2951 |
| Negative | 175-250 ms | 12.5-16.5 Hz | 10 | -2.9483 | -4.5112 | 0.3665 |
| Negative | 0-100 ms | 10.8-16.0 Hz | 13 | -2.5266 | -3.5713 | 0.4287 |
| Negative | 525-600 ms | 10.2-14.2 Hz | 6 | -2.5424 | -3.4094 | 0.5283 |
| Negative | 650-700 ms | 12.8-16.0 Hz | 4 | -2.5818 | -3.4690 | 0.6479 |
| Negative | 650-700 ms | 18.5-21.5 Hz | 5 | -2.4293 | -3.1452 | 0.7149 |
| Negative | 950-1000 ms | 15.2-18.5 Hz | 7 | -2.5502 | -3.1816 | 0.8108 |
| Negative | 625-650 ms | 17.5-20.2 Hz | 4 | -2.4452 | -3.2788 | 0.8294 |
| Negative | 125-150 ms | 20.8-24.0 Hz | 3 | -2.5469 | -3.3648 | 0.8500 |
| Negative | 425-475 ms | 5.8-7.0 Hz | 4 | -2.7347 | -3.6687 | 0.8540 |
| Negative | 175-200 ms | 12.8-15.0 Hz | 3 | -2.7301 | -3.5709 | 0.8572 |
| Negative | 725 ms | 16.2-19.5 Hz | 4 | -2.4088 | -2.6793 | 0.8610 |
| Negative | 225-250 ms | 23.8-25.5 Hz | 3 | -2.6735 | -3.8220 | 0.8934 |
| Negative | 1000 ms | 15.5-17.2 Hz | 4 | -2.3040 | -2.4428 | 0.9264 |
| Negative | 0-25 ms | 9.5-10.5 Hz | 3 | -2.3772 | -2.7550 | 0.9460 |
| Negative | 750 ms | 18.8-19.5 Hz | 4 | -2.2833 | -2.7353 | 0.9550 |
| Negative | 425 ms | 18.5-19.5 Hz | 3 | -2.3967 | -2.8655 | 0.9560 |
| Negative | 275 ms | 12.2-12.5 Hz | 5 | -2.3842 | -2.8644 | 0.9646 |
| Negative | 975 ms | 15.0-15.2 Hz | 5 | -2.2476 | -2.5234 | 0.9646 |
| Negative | 650 ms | 17.8-18.2 Hz | 3 | -2.3927 | -2.9532 | 0.9652 |
| Negative | 250 ms | 12.2-12.5 Hz | 4 | -2.4117 | -2.8403 | 0.9694 |
| Negative | 75 ms | 17.5-17.8 Hz | 4 | -2.3105 | -2.6264 | 0.9696 |
| Negative | 325 ms | 8.0-8.2 Hz | 4 | -2.2558 | -2.5260 | 0.9708 |
| Negative | 950 ms | 23.8-24.0 Hz | 3 | -2.1760 | -2.3234 | 0.9734 |
| Negative | 275 ms | 12.0 Hz | 4 | -2.3218 | -2.7453 | 0.9758 |

**Supplementary Table S3**: The cluster-based and target time-locked permutation test results were analyzed from 0 to 1000 ms and 4 to 30 Hz. One significant negative cluster was found; the asterisk represents the significant results (p < .05*).

#### *3.2 Significant TF results in post-target window*

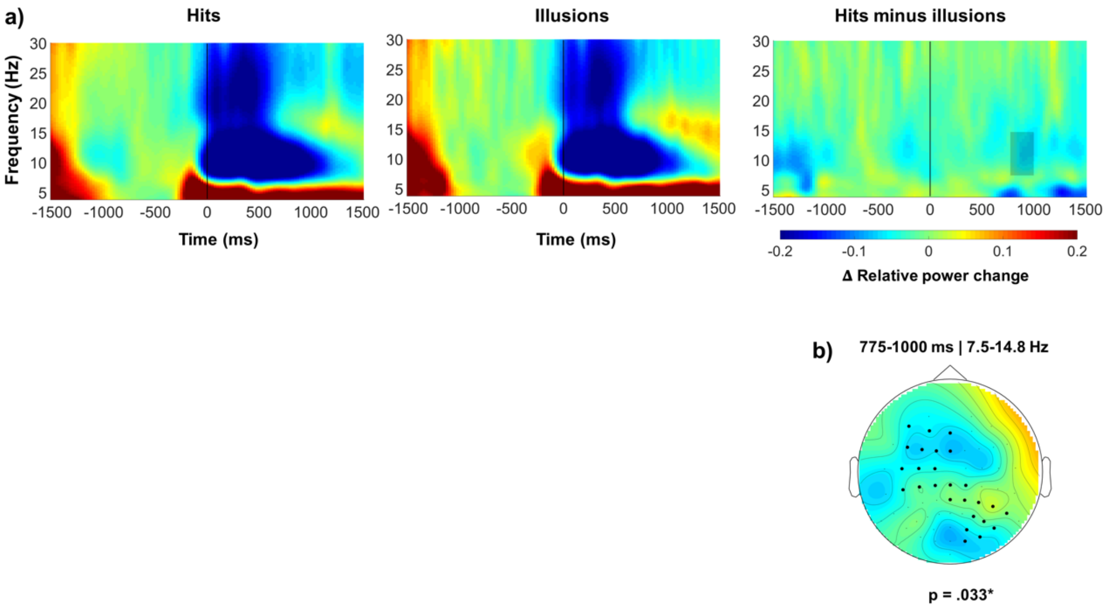


**Supplementary Figure S3**: Cluster-based permutation analysis representation across the completed frequency range (4-30 Hz). The analyzed window was from 0 to 1000 ms after target onset. a) The figure shows the TF map for hits, illusions, and the difference between hits and illusions (from left to right). The shaded area in the conditions difference TF map represents the significant time interval (775-1000 ms) and frequency range (7.5-14.8 Hz). b) Topographic distributions of the averaged t-values within the significant time-frequency intervals; black dots indicate electrodes contributing to the cluster-level statistic; the color bar is the same for the TF map.

### 4. Additional control analysis

#### *4.1 R-peak timing relative to target onset*

To verify that heartbeat-locked effects were not confounded by differences in the temporal relationship between selected R-peaks and target onset, we compared the mean R-peak-to-target interval between hits and illusions. By design, selected R-peaks occurred between 1600 and 600 ms before target onset, and heartbeat-locked analyses were restricted to the 200-600 ms post-R-peak interval. Therefore, the analysed window was always strictly pre-target and could not overlap with target-evoked activity.

A paired-samples t-test revealed no significant difference in the temporal distance between selected R-peaks and target onset for hits and illusions, $t(24) = 0.28$, $p = .782$, Cohen's $d_z = 0.06$ (see Figure S4). These results indicate that heartbeat timing relative to target presentation was comparable across conditions and therefore cannot account for the observed HEP or HEO differences.

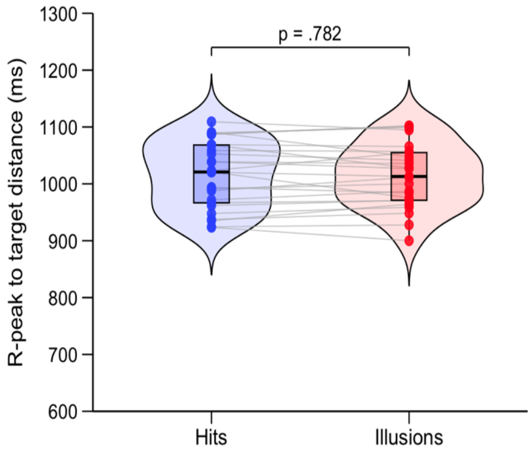


**Supplementary Figure S4**: R-peak timing relative to target onset for hits and illusions. Violin plots show the distribution of participant-level mean R-peak-to-target intervals for each condition. Boxplots indicate the median and interquartile range, dots represent individual participants, and grey lines connect paired observations. No significant difference was observed between hits and illusions, $t(24) = 0.28$, $p = .782$.

#### *4.2 HEP Surrogate R-peak*

To assess whether the reported HEP effect depended on the temporal correspondence between heartbeat timing and neural activity, a surrogate-heartbeat control analysis was performed following the procedure described by Steinfath et al. (2025). For each participant and condition, heartbeat timing was randomly reassigned across trials while preserving the original EEG data and condition labels. This procedure disrupts the temporal correspondence between cardiac events and neural activity while preserving any condition-related EEG differences unrelated to heartbeat timing.

A total of 1,000 surrogate datasets were generated. For each surrogate dataset, the same cluster-based permutation procedure used in the original HEP analysis was repeated. The maximum absolute cluster statistic across positive and negative clusters was retained from each iteration, yielding a two-sided surrogate distribution of cluster statistics.

The original HEP cluster had an absolute cluster statistic of $|sum(t)| = 488.899$, exceeding the 95th percentile of the surrogate distribution (455.739), with an empirical two-sided $p = .042$. (see Figure S5) Thus, effects of the magnitude observed in the original analysis occurred infrequently after disruption of heartbeat–EEG alignment. These findings suggest that the HEP effect depended on the temporal alignment between heartbeat timing and neural activity and cannot be fully explained by condition-related EEG differences independent of cardiac timing.

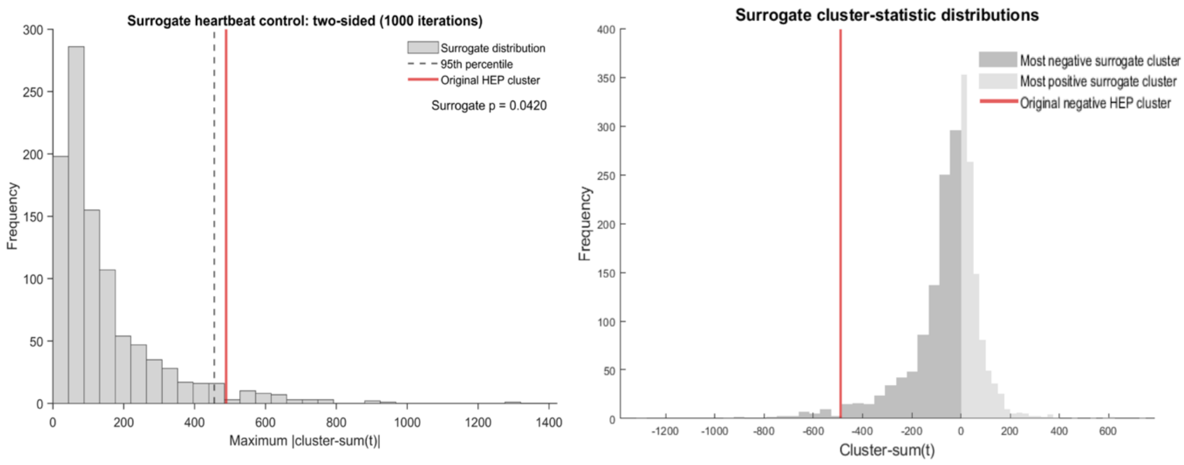


**Supplementary Figure S5**. Surrogate heartbeat control of the HEP effect. Left panel: Two-sided surrogate distribution of the maximum absolute cluster statistic across 1,000 iterations. The red line indicates the original HEP cluster and the dashed line indicates the 95th percentile of the surrogate distribution (empirical p = .042). Right panel: Signed distributions of the most negative and most positive surrogate cluster statistics; the red line indicates the original negative HEP cluster. Inferential testing was based on the two-sided maximum absolute cluster-statistic distribution shown in the left panel.

#### *4.3 HEO Surrogate R-peak*

The same surrogate-heartbeat procedure described in Section 4.2 was applied to the HEO analysis. Due to the substantially greater computational demands of the time-frequency analyses, 300 surrogate datasets were generated, and the number of randomisations in the cluster-based permutation procedure was reduced from 5,000 to 1,000. This modification preserved the inferential framework of the analysis while reducing computational time.

The original HEO cluster had an absolute cluster statistic of |sum(t)| = 4,987.030, exceeding the 95th percentile of the surrogate distribution (3,847.664), with an empirical two-sided p = .035 (see Figure S6). Thus, effects of this magnitude occurred infrequently after disruption of heartbeat–EEG alignment. These findings suggest that the observed HEO effect depended on the temporal alignment between heartbeat timing and ongoing neural activity and cannot be fully explained by condition-related prestimulus EEG differences independent of cardiac timing.

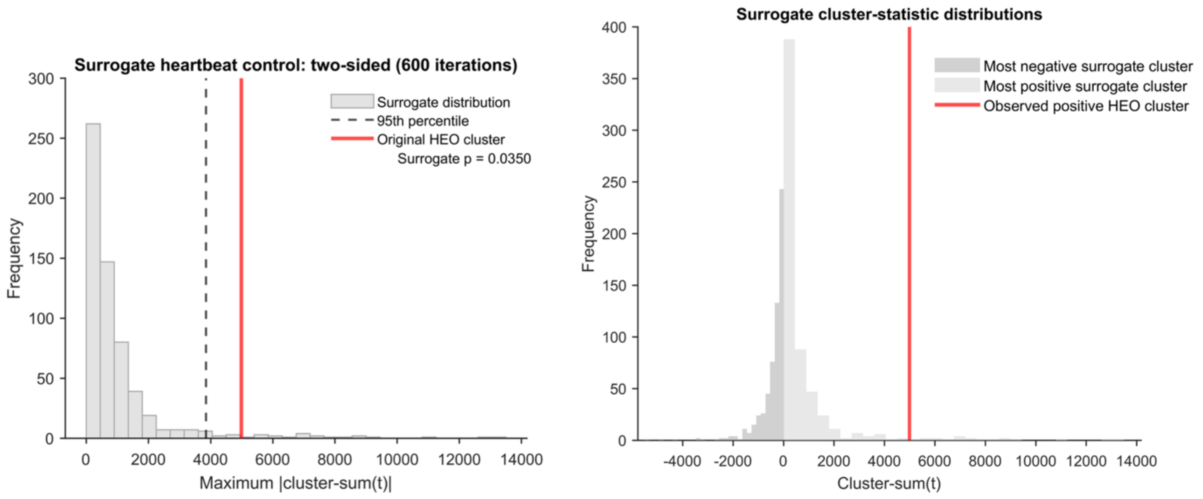


**Supplementary Figure S6**. Surrogate heartbeat control of the HEO effect. Left panel: Two-sided surrogate distribution of the maximum absolute cluster statistic across 300 iterations. The red line indicates the original HEO cluster and the dashed line indicates the 95th percentile of the surrogate distribution (empirical p = .035). Right panel: Signed distributions of the most negative and most positive surrogate cluster statistics; the red line indicates the original positive HEO cluster. Inferential testing was based on the two-sided maximum absolute cluster-statistic distribution shown in the left panel.

### 5. Trial-level analysis

#### *5.1 HEP amplitude and perceptual outcome*

To assess whether trial-to-trial variations in prestimulus HEP amplitude were associated with subsequent perceptual outcome, we conducted exploratory single-trial analyses using generalized linear mixed-effects models (GLMMs) with a binomial link function implemented in lme4 (Bates et al., 2015). To reduce circularity in feature extraction, HEP amplitude was quantified using a leave-one-subject-out (LOSO) procedure. For each participant, amplitude estimates were extracted from a participant-specific channel × time mask derived from the remaining participants. HEP amplitude and R-peak-to-target latency were decomposed into within- and between-participant components using participant-mean centering (Enders & Tofighi, 2007; Curran & Bauer, 2011), standardized, and entered into a GLMM with random intercepts for participant.

The analysis included 9,221 trials from 25 participants. More negative HEP amplitudes relative to a participant's own mean were associated with a lower probability of reporting an illusion (HEPWithin: β = −0.078, OR = 0.925, 95% CI [0.884, 0.968], p = .00074; see Figure S7). Expressed differently, a one-standard-deviation increase in HEP negativity corresponded to approximately 7.5% lower odds of an illusion. No significant between-participant effect was observed (p = .393), indicating that the association was driven by trial-to-trial fluctuations within individuals rather than by stable differences in HEP amplitude between participants.

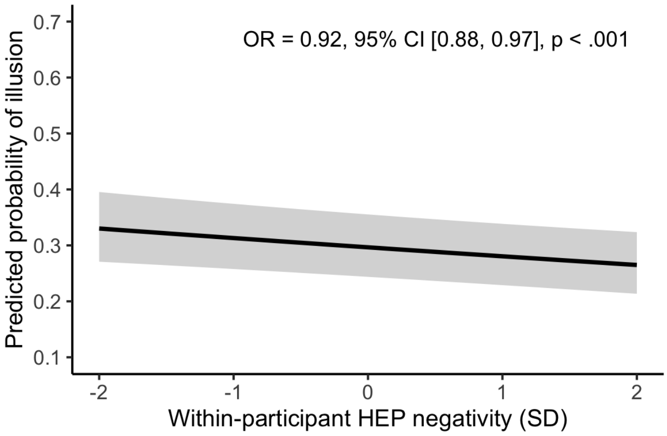


**Supplementary Figure S7**: Trial-level HEP amplitude and perceptual outcome. Predicted probability of reporting an illusion as a function of within-participant HEP amplitude estimated from the generalized linear mixed-effects model. Greater HEP negativity was associated with a lower probability of reporting an illusion. The shaded area represents the 95% confidence interval.

R-peak-to-target latency was not associated with perceptual outcome at either the within-participant ($p = .863$) or between-participant ($p = .444$) level, suggesting that the observed relationship was not explained by variation in heartbeat timing relative to target onset. Model diagnostics indicated no evidence of over- or underdispersion (dispersion = 0.998, $p = .944$), and multicollinearity was negligible (all VIFs ≤ 1.18).

Overall, the results suggest that trial-to-trial fluctuations in prestimulus HEP amplitude may be associated with subsequent perceptual outcome.

*5.2 HEO power and perceptual outcome*

To investigate whether trial-to-trial fluctuations in HEO were associated with subsequent perceptual outcome, an analogous LOSO-based GLMM approach described for the HEP analysis was applied to the alpha, early-beta, and late-beta HEO effects identified in the primary time-frequency analyses.

The analyses included 7,882 valid trials from 25 participants. Across all three frequency bands, greater prestimulus HEO power was associated with lower odds of reporting an illusion, consistent with the direction of the condition-level findings. However, none of the within-participant effects reached statistical significance following LOSO-based feature extraction: AlphaWithin ($\beta = -0.042$, OR = 0.959, 95% CI [0.911, 1.010], $p = .114$), BetaEarlyWithin ($\beta = -0.037$, OR = 0.964, 95% CI [0.917, 1.014], $p = .155$), and BetaLateWithin ($\beta = -0.030$, OR = 0.970, 95% CI [0.923, 1.020], $p = .238$). No significant between-participant effects were observed (all ps > .093).

A joint model simultaneously including alpha, early-beta, and late-beta predictors likewise revealed no significant independent contributions of any oscillatory measure to perceptual outcome (all ps > .22). Multicollinearity was low (all VIFs ≤ 4.06), and simulation-based residual diagnostics showed no evidence of over- or underdispersion (all dispersion values ≈ 1.00; all ps ≥ .958).

Taken together, the trial-level HEO analyses showed effects in the same direction as the group-level findings, with greater alpha and beta power generally associated with a lower probability of reporting an illusion. However, these associations did not reach conventional levels of statistical significance following independent LOSO-based feature extraction.

### 6. Supplementary top-down expectancy analysis, results and discussion

#### *6.1 Analysis plan*

To assess the role of top-down expectancy, the analytical pipelines described in Sections 2.3.1 (HR), 2.3.2 (HEP), 2.3.3 (HEO), and 3.5 (correlations) were extended to include the Awareness factor (aware vs. unaware) as a between-subjects factor. For HR, this involved incorporating Awareness and its interactions with Trial type and Time within the LME fixed-effects. For HEP and HEO, the same cluster-based permutation approach was applied to compare aware and unaware participants using an independent-samples t-test as the sample-level statistic. Finally, the same correlation pipeline was applied to examine effects associated with the Awareness factor.

#### *6.2 HR, HEP, HEO and correlation results*

When the Awareness factor was added to the HR analysis, no main effect reached significance ($X^2(1) = 0.1264$; $p=.722$; $\eta^2 = 8.22^{-3}$). Because Awareness did not explain additional variance at the main level, interactions involving this factor were not further examined. In the HEP data, analyses conducted separately within the aware and unaware groups did not reveal differences between hits and illusions in the early (all ps > .438) or late (all ps > .106) window in either group. Likewise, no differences were observed when directly comparing HEP amplitude between aware and unaware participants, irrespective of trial type (all ps > .735). For the HEO results, the between-groups contrast for awareness did not reveal any significant effects (all ps>.418) across the entire range of frequencies. Regarding correlations, the same analytical pipeline in section 3.5 in the main manuscript was applied to the between-group factor Awareness, but none of the effects reached significance (all ps> .239).

#### *6.3 Discussion*

Against our predictions, no effects of top-down expectations were observed. Although previous studies in our lab reported differences in gamma power (Cobos et al., 2023) and HR (Cobos et al., 2026) between aware and unaware participants, such effects were not found in the present study. Importantly, null or limited effects of expectation have been reported, particularly at early perceptual stages or when expectations do not directly influence sensory encoding (Rungratsameetaweemana et al., 2018; Zivony & Eimer, 2024). More broadly, the literature shows mixed evidence regarding whether expectations consistently modulate sensory representations (Zhou et al., 2020).

From the perspective of the binding framework proposed by Scholte & De Haan (2025), feature integration failures are not conceived as the breakdown of a specialized binding mechanism but rather as the consequence of increased ambiguity when strongly learned feature co-occurrences are violated. In the present study, participants were initially trained to integrate a specific stimulus configuration, but violating expectations toward the end of the task increased perceptual ambiguity, such that some participants became aware of the change while others did not. We hypothesized that cardiac-brain coupling, indexed by HEP and HEO, would differentially reflect this manipulation of expectations; however, no significant effects emerged. This null result may be partly attributable to limited statistical power within the aware and unaware subgroups. Importantly, these findings do not rule out a role for cardiac signals in resolving perceptual ambiguity but rather highlight the need for future studies with larger samples and targeted manipulations to further investigate how interoceptive signals contribute to whether expectancy violations reach awareness under ambiguous conditions. Accordingly, a larger sample size may be required to effectively compare aware and unaware participants, as the current subgroups (n = 11 and n = 14) are statistically underpowered to detect subtle differences (Baker et al., 2021; Szucs & Ioannidis, 2017).
